# Early life stress exposure alters brain vasculature transcriptomic profiles in areas regulating stress resilience

**DOI:** 10.64898/2026.04.16.718991

**Authors:** Jose L. Solano, Béatrice Daigle, Manon Lebel, Catherine Jensen Peña, Caroline Ménard

## Abstract

Early life stress (ELS) events during sensitive postnatal time periods can recalibrate future stress responsiveness and precipitate mental disorders. Neurovascular adaptations can influence cognition, mood, and stress responses. Disruption of blood-brain barrier (BBB) integrity, which is formed by endothelial cells, astrocytes, and pericytes, has been implicated in affective disorders such as depression, which often arise from chronic stress experiences. Despite the BBB undergoing critical maturation stages during development, it remains poorly known how ELS influences brain vascular function, as previously shown for adult stress, and whether it augments BBB vulnerability to subsequent challenges. First, we took advantage of a public two-hit stress RNA-sequencing dataset and filtered for vascular enriched genes in the prefrontal cortex and nucleus accumbens, the two brain regions where BBB integrity is frequently compromised. This analysis revealed BBB-related gene ontology categories modulated by either ELS alone or its combination with adult stress. Then, using a mouse model combining ELS with chronic social defeat stress (CSDS) in adulthood, we found that ELS did not exacerbate CSDS susceptibility; instead, it increased social interactions and the likelihood of a resilient profile in both males and females. Transcriptomic profiling in our cohort further identified distinct sex- and region-specific BBB gene expression patterns associated with ELS and its interaction with CSDS. Additionally, we observed a reduction of corticosterone levels, the primary stress hormone, following CSDS. Altogether, these results indicate that ELS modulates stress responses when facing emotional challenges in adulthood, possibly through long-lasting changes of BBB function via the glucocorticoid system.

**Highlights:**

- RNA-seq vascular filtering reveals BBB distinct ontology categories for ELS and AS
- ELS increases the likelihood of a high social and resilient profile.
- Pericytes gene expression associated to resilience is sex- and region-specific.
- CORT response desensitizes after adult CSDS in both sexes.

## 1. Introduction

Converging evidence from preclinical, clinical, and epidemiological studies indicates that early-life stress (ELS) can, under specific conditions, calibrate future stress responsiveness in an adaptative manner promoting resilience (Hartmann and Schmidt, 2020; Sisk et al., 2025; Torres-Berrío et al., 2026). For example, in rodent models, brief and predictable stressors, like short periods of maternal separation paired with supportive rearing environments, have been shown to enhance stress-coping abilities and improve emotional regulation later in life (Santarelli et al., 2017; Shi et al., 2021). Even though context-dependent adaptations are linked to ELS positive outcomes, ELS exposures are frequently detrimental and constitute a risk factor for development of mental and physical health problems (Bonapersona et al., 2019; Duffy et al., 2018; Koolhaas et al., 2011; Nelson et al., 2020). The type, the timing, and the cumulative load of stressful experiences such as physical or emotional abuse, parental neglect, or prolonged unpredictable maternal separation in rodents, strongly predict harmful cognitive, behavioral, and emotional outcomes (Nelson et al., 2020; Duffy et al., 2018; Koolhaas et al., 2011). Mechanisms through which ELS disrupts neurodevelopment, alters Hypothalamic-Pituitary-Adrenal (HPA) axis signaling, and sensitizes immune responses have been extensively documented (e.g., Dutcher et al., 2020; Malave et al., 2022; Murthy and Gould, 2018; Nelson et al., 2020; Parise et al., 2025). Importantly, ELS also induces adaptations beyond the central nervous system, impacting systemic and peripheral physiology. In both humans and rodents, ELS has been linked to chronic inflammation, altered immune-vascular communication, and increased cardiometabolic risk, emphasizing whole-body interactions that extend beyond neural circuits (Doney et al., 2022; Dutcher et al., 2020; Kellum et al., 2025).

The blood–brain barrier (BBB) is a dynamic neurovascular interface formed by endothelial cells, pericytes, and astrocytes maintaining CNS homeostasis while adapting to environmental challenges and protecting the brain from deleterious signals circulating in the blood (Chagnot and Montagne, 2025; Dion-Albert et al., 2023). In adulthood, chronic social defeat stress (CSDS) induces BBB hyperpermeability via loss of tight junction protein Claudin-5 (Cldn5) in a sex- and brain region-specific manner with a higher vulnerability in the nucleus accumbens (NAc) for males vs prefrontal cortex (PFC) for females (Dion-Albert et al., 2022b; Dudek et al., 2020; Menard et al., 2017; Paton et al., 2026). Disruption of the BBB within these regions, implicated in mood regulation, reward and social behavior, has been causally linked to stress vulnerability, social avoidance and depressive-like behaviors, whereas intact BBB function and beneficial adaptations are associated with stress resilience (Dion-Albert et al., 2022a; Dudek et al., 2025; Paton et al., 2023). Even though most of the studies demonstrating how BBB integrity is compromised after stress used adult paradigms, in rodents, it has been shown that BBB permeability can be increased when either maternal separation (Gómez-González and Escobar, 2009) and limited bedding and nesting (Evertse et al., 2026) are implemented during the first postnatal weeks. Interestingly, when maternal separation is followed by an immune challenge at weaning or later in adult life, ELS persistently altered the expression of endothelial tight junctions (e.g., Cldn5 and Occludin) (Solarz et al., 2023). Taken together, this evidence indicates that ELS can heighten vascular vulnerability to stress. However, it remains unclear whether the BBB could encode an ELS challenge, for example via alterations in the transcriptomic profiles of vascular cells, leading to altered responses to subsequent challenges.

Here we leveraged a two-hit stress paradigm to address this knowledge gap and tested how the BBB adapts to stress across life when facing an initial exposure during childhood followed by another one in adulthood. First, we explored a public RNA-sequencing (RNA-seq) dataset from a study using a two-hit stress (Peña et al. 2019, 2017) and filtered for genes enriched in brain vascular cells to identify BBB-related gene ontology categories in the PFC and NAc. Enrichment for pathways related to *blood vessels morphogenesis, blood vessel development* and *angiogenesis* guided our hypothesis-driven transcriptomic assays. Then, we generated our own dataset by implementing a similar two-hit stress model, maternal separation combined with limited bedding and nesting as ELS followed by adult CSDS, in both sexes. Behavioral assessment before and after CSDS challenged our initial hypothesis of ELS-induced exacerbated stress vulnerability with increased sociability and stress resilience after ELS. Accordingly, analysis of the glucocorticoid (GC) system revealed a desensitized corticosterone (CORT) response consistent with a stress-inoculation framework driving adult resilience (Crofton et al., 2015; Hartmann and Schmidt, 2020). Profiling of BBB gene expression further revealed that ELS predominantly modulated pericyte-associated genes in the male NAc and female PFC, and astrocyte-associated genes in the male NAc, suggesting that those cells might be neurovascular drivers of resilience in a sex- and brain region-specific manner.

## 2. Material and methods

### 2.1. Animals

Male and female C57BL/6 mice of 7-8 weeks of age were purchased from Charles River for breeding at the CERVO Brain Research Center (Quebec City, Canada). After one week of acclimatation, reproduction was initiated by pairing one male with two females. For this study, 122 mice (63 males/59 females) born at CERVO from primigravida mothers were included. Weaning was set at P21 with males and females separated by sex (2-4 mice/cage). Littermates were kept together and pups from the same experimental condition combined if necessary to avoid isolation. Standard housing conditions consisted of regular bedding, 1 cotton nesting square, a plastic house and a plastic chew toy. For the CSDS paradigm, retired male CD-1 breeders from Charles River (~40 g) of at least 4 months of age were used as aggressors (AGG). CD-1 retired breeders were single housed with access to bedding for the duration of the social defeat. For all animals, room temperature was maintained between 19 and 23 °C, with a 12 h/12 h light/dark cycle and humidity was kept around 40−45%. Water and food were provided ad libitum. All mouse procedures were performed in accordance with the Canadian Council on Animal Care (1993) as well as Université Laval animal care committee (Certificate #2021-937).

### 2.2. Stress interventions

#### 2.2.1. Early-Life Stress

Early-life stress (ELS) protocol was a combination of maternal separation and limited bedding and nesting, as previously described by Peña et al. (2017). Animals were assigned randomly to the control (ELS-) or to the ELS (+) group. For ELS+, litters were separated from their mother 4h per day for 10 days, from post-natal day 10 (PD10) to PD19. In addition, during the same period, bedding and nesting was reduced to 1/3 of the standard conditions. For ELS-litters, the plastic chew toy and the plastic house were removed, and a short handling was performed each day. At P19 after the last separation cycle, environmental conditions were restored to standard housing conditions and maintained until adulthood.

#### 2.2.2. Adult Chronic Stress

Mice were exposed to 10days of CSDS from PD61 to PD70. Prior defeat, CD-1 mice were screened for aggressive behaviour by assessing inter-male social interactions for three consecutive days and then housed in the social defeat cage (26.7 cm width × 48.3 cm depth × 15.2 cm height, Allentown Inc) 24 h before the beginning of CSDS. For males, the paradigm was performed as detailed in Golden et al. (2011) and Menard et al. (2017). For females, aggression bouts were triggered by applying CD-1 male urine as previously described by Harris et al. (2018) and Dion-Albert et al. (2022). Each C57BL/6 female mouse was paired with the urine from a particular CD-1 aliquot (BioIVT MSE00URINE-0110414) throughout the entire course of CSDS. Each day, before physical interaction with an unfamiliar CD-1 aggressor, urine was applied (20 ul/area) to the base of the tail, vaginal orifice and upper back of the female mouse. Physical interactions with a CD-1 aggressor lasted for 5 minutes (males) up to 10 minutes (females). After the antagonistic encounter, the male or female C57BL/6 mouse were housed on the opposite side of the social defeat cage divided by a clear perforated Plexiglas divider (0.6 cm × 45.7 cm × 15.2 cm), allowing sensory contact, over the subsequent 24h. This was repeated for 10 days with a novel CD1 mouse each day. Throughout the sessions, mice were monitored for aggressive interactions and mounting behaviours. A session was immediately stopped if persistent mounting or fighting causing physical wounding occurred. Unstressed control mice (CSDS-) were housed in similar defeat cages two per cage on either side of a perforated divider and rotated daily. Both CSDS- and CSDS+ mice were singly housed after the last bout of defeat.

### 2.3. Behavioral test

All tests were recorded and monitored using the video tracker software EthoVision XT 17 (Noldus). Locomotion and cumulative time in zones were automatically calculated and used for statistical analyzes and group comparisons. Social and anxiety-like behaviors were evaluated before and after CSDS. The open field test (OF) and the social preference test (SPT) were implemented before CSDS, while the social interaction test (SI) and the elevated plus maze were implemented after CSDS (**Fig.2A**).

#### 2.3.1. Open Field Test

Locomotor activity was first assessed at PD56 during 10 min in a squared white acrylic arena (50 width x 50 depth x 50 height cm). The OF arena was uniformly illuminated from the ceiling with white light. To evaluate anxiety-like behaviors, the OF was divided into two zones for posterior analysis: center and periphery (Novoa et al., 2022; Seibenhener and Wooten, 2015). Finally, the same OF was used as habituation period for the social preference test (SPT).

#### 2.3.2. Social Preference Test

Twenty-four hours after OF, a SPT was performed to assess sociability. The SPT test consisting in 2 trials of 5 min, was performed under white light conditions. In trial 1, two small wire cages were placed in opposite corners of the OF arena. It allowed to evaluate novelty-behavior, baseline exploratory behaviour, locomotion and the natural “biased” preferred zone for each animal (Netser et al., 2017; Rein et al., 2020). For trial 2, a social target (a novel mouse matching strain, age and sex) was placed inside the wire cage in the zone were the animal spent the less time during the previous trial, thus limiting place preference bias. The SPT arena was divided in four zones: two empty areas (opposing corners), 1 including an empty cage and 1 with either an empty cage or social target. A social preference score was calculated as follows: *(Time with social target - Time with empty cage) / (Time with social target + Time with empty cage)*. In addition, a SPT ratio was calculated as follows *Time with social target / Time with preferred empty cage in trial 1*. Animals were categorized as highly social when reaching the following criteria: SPT ratio “>1” and time spent with the social target “> 50%” during trial 2. Low sociability was defined as: SPT ratio “<1” and time spent with the social target “< 50%” of the time during trial 2. If animals had an ambiguous score, they were classified as indifferent.

#### 2.3.3. Social Interaction Test

24h after the last defeat bout, a SI test consisting of 2 trials of 2.5 min was performed under red-light conditions as previously described (Golden et al., 2011; Menard et al., 2017). For this test, the same arena previously described was used. First, the C57BL/6 male or female mouse was allowed to freely explore the arena with an empty large wire animal cage placed at one end. This trial was used to determine baseline exploratory behaviour and locomotion in the absence of a social target. Next, a novel aggressive male CD-1 (AGG) was introduced, and exploratory behavior was again evaluated. The SI ratio was calculated as follows *Time in the interaction zone with the AGG present/absent*. Animals were categorized as highly social when the SI ratio was “>1” and time in the interaction zone spent with the AGG “> 50%”. Low sociability was defined as SI ratio “<1” and time in the interaction zone spent with the AGG “< 50%”. If animals had an ambiguous score, they were classified as indifferent. Based on the classical SI ratio cut-off following CSDS exposure (Golden et al., 2011), highly social animals were also considered resilient (RES) while mice with low sociability were labelled as stress-susceptible (SUS). Finally, to unravel biological mechanisms driving RES vs SUS, indifferent animals were removed of CORT and transcriptomic analysis (Peña et al., 2019b, 2017).

#### 2.3.4. Elevated Plus Maze

4h after the SI, animals were placed in the middle of a white Plexiglas cross-shaped elevated plus maze (EPM) under white light conditions for 5 min. The maze consists of a center area, two open arms (arms of 12 cm width × 50 cm length) without walls and two black closed arms (40 cm high) set on a pedestal 1 m above floor level. This test evaluates anxiety-like behavior by modulation of the natural aversion of rodents to open and illuminated areas (Walf and Frye, 2007).

### 2.4. qPCR

At PD72, 24h after the SI and EPM, bilateral 2.0 mm brain punches of the prefrontal cortex (PFC) and nucleus accumbens (NAc) were collected from 1-mm coronal slices and immediately flash frozen and stored at −80°C. RNA was isolated with TRIzol (Invitrogen) homogenization and chloroform layer separation using the Pure Link RNA mini kit (Life Technologies). RNA concentration was determined on a Take3 microvolume plate reader (EON 12120514) and reversed transcribed to cDNA with the Maxima-H-minus cDNA synthesis kit (Life Technologies). Each qPCR reaction contained 3 ng of sample cDNA, 5 μL of Power up SYBR green (Life Technologies), 1 μL of PrimeTime qPCR primer (Integrated DNA Technologies) and 1 μL ddH2O. Samples were heated at 95°C for 2 mins, followed by 40 cycles of 95°C for 15 s, 60°C for 33 s and 72°C for 33 s. The ΔΔCt method was used for analysis, with normalization on mouse Gapdh, a housekeeping gene. Primer pairs are listed in **Supp.Table1**.

### 2.5. Corticosterone ELISA

Blood samples were collected at PD58 via submandibular vein puncture and at PD72 after rapid decapitation for brain collection. Blood was allowed to clot for at least 1 h before being centrifuged at 10,000 RPM at room temperature for 2 min. The supernatant was collected and spin again at 3000 RPM for 10 min. The clear supernatant (serum) was collected, aliquoted and stored at −80°C until use. To determine CORT levels, serum samples were processed using the DetectX^®^ ELISA kit (Arbor Assays # K014-H, sensitivity 14.35pg/mL and assay range 19.53pg/mL to 5000pg/mL) at 1:100 dilution as recommended by the manufacturer. Optical density was measured at 450 nm with a plate reader (EON 12120514) and concentrations calculated from a 4-PL standard curve. Samples with a coefficient of variation above 15% were removed from the analysis.

### 2.6. Public bulk RNA-Seq analysis for neurovascular cells

To evaluate the plausible impact of two-hits of stress on the BBB-associated transcriptomic profile, an independent and unbiased genome-wide data set from Peña et al. (2019, 2017), GSE89692, https://www.ncbi.nlm.nih.gov/geo/query/acc.cgi?acc=GSE89692) was filtered considering genes enriched in brain vascular cells identified by Valandewijck et al. (2018) (**Supp.Table2**). Then, the impact of combining ELS and adult chronic stress (AS, CSDS for males and subchronic variables stress for females) on BBB genes was assessed by performing DESeq2 on the 3315 filtered genes. Main effects of both ELS and AS were evaluated with the likelihood ratio test. Differentially enriched genes (DEG) were used to identified enriched ontology terms (clusterProfiler::enrichGO).

### 2.7. Statistical analysis

Data analysis was performed using R version 4.4.3 and Rstudio 418. The list of associated packages is listed in **Supp.Table3**. Parametric and non-parametric (permutation tests) two-way ANOVAs followed by Benjamini–Hochberg’s post hoc were performed to determinate sex, ELS and CSDS effects. Parametric or non-parametric analysis were performed when appropriate considering normality (Shapiro-Wilk test) and homoscedasticity (Levene test) assumptions. If normality was met but not homoscedasticity, the Welch adjustment was applied. Multinominal modeling was performed to determine the odds of high sociability for the SPT and SI test. To identify the proportion of RES and SUS after CSDS, a binomial model was applied. For gene expression, analyses were performed within each sex groups, and relative gene expression was estimated using ELS-:CSDS-as reference, for both hierarchical clustering and group comparison. Spearman correlations were performed between serum CORT levels and expression of GC-related genes. Significance thresholds were set at p<0.05 with tendencies at p<0.07.

## 3. Results

### 3.1. Two-hits of stress impact the blood-brain barrier transcriptomic

ELS and AS are two well characterized stressors representing risk factors for the development of cardiovascular (Kellum et al., 2025; Kelly et al., 2023) and mental health-related conditions (Dion-Albert et al., 2022a; Duffy et al., 2018). They impact several biological domains, including but not limited to epigenetic, immune and neuronal systems, both in human and animal models (Malave et al., 2022; Nelson et al., 2020). Even further, their combination generally has summatory effects with detrimental consequences on anxiety- and depression-like behavior (Bonapersona et al., 2019; Peña et al., 2019a; Rincel et al., 2019). Recent evidence suggest that ELS can disrupt BBB tight junctions in the striatum inducing impairment in social behaviors and effort-based reward learning in adulthood (Evertse et al., 2026). However, the impact of two-hits of stress on the brain vascular transcriptional profiles remain to be clarified. To address this knowledge gap, we first took advantage of a public bulk RNA-sequencing dataset (Peña et al., 2019b, 2017) and filtered genes relevant to cells of the perivascular space with an adult mouse brain vascular atlas (Vanlandewijck et al., 2018). We chose to focus on PFC and NAc which our group and others have reported stress-inducing loss of BBB integrity in these brain regions for females and males, respectively (Dion-Albert et al., 2022b; Dudek et al., 2020; Gal et al., 2023; Menard et al., 2017; Shi et al., 2024). Intriguingly, ELS combined with AS (ELS + AS) modified directionality (up and down) BBB transcriptional changes when compared to ELS or AS alone (**Fig.1A,C,E**). In the male PFC, stress exposure regulated 129 common genes and the combination of ELS + AS differentially altered 64 genes (**Fig.1A**). These differentially expressed genes (DEGs) were tied to gene ontology (GO) terms like *blood vessel development, blood vessel morphogenesis, angiogenesis*, and *cell proliferation processes* (**Fig.1B, Supp.Table4**). Meanwhile, in the NAc, no meaningful change was noted. For females, stress exposure regulated 52 common genes in the PFC vs. 72 genes in the NAc, and the combination of ELS + AS differentially regulated 67 genes for the PFC and 72 genes for the NAc (**Fig.1B,E**). Female PFC DEGs were associated with modulation of the *extracellular matrix, cell migration, blood vessel development*, and *blood vessel morphogenesis* (**Fig.1D, Supp.Table5**). As for NAc DEGs, GO terms were linked to *vesicles membrane, anchoring junction*, and *protein localization* (**Fig.1F, Supp.Table6**).

**Fig.1.**
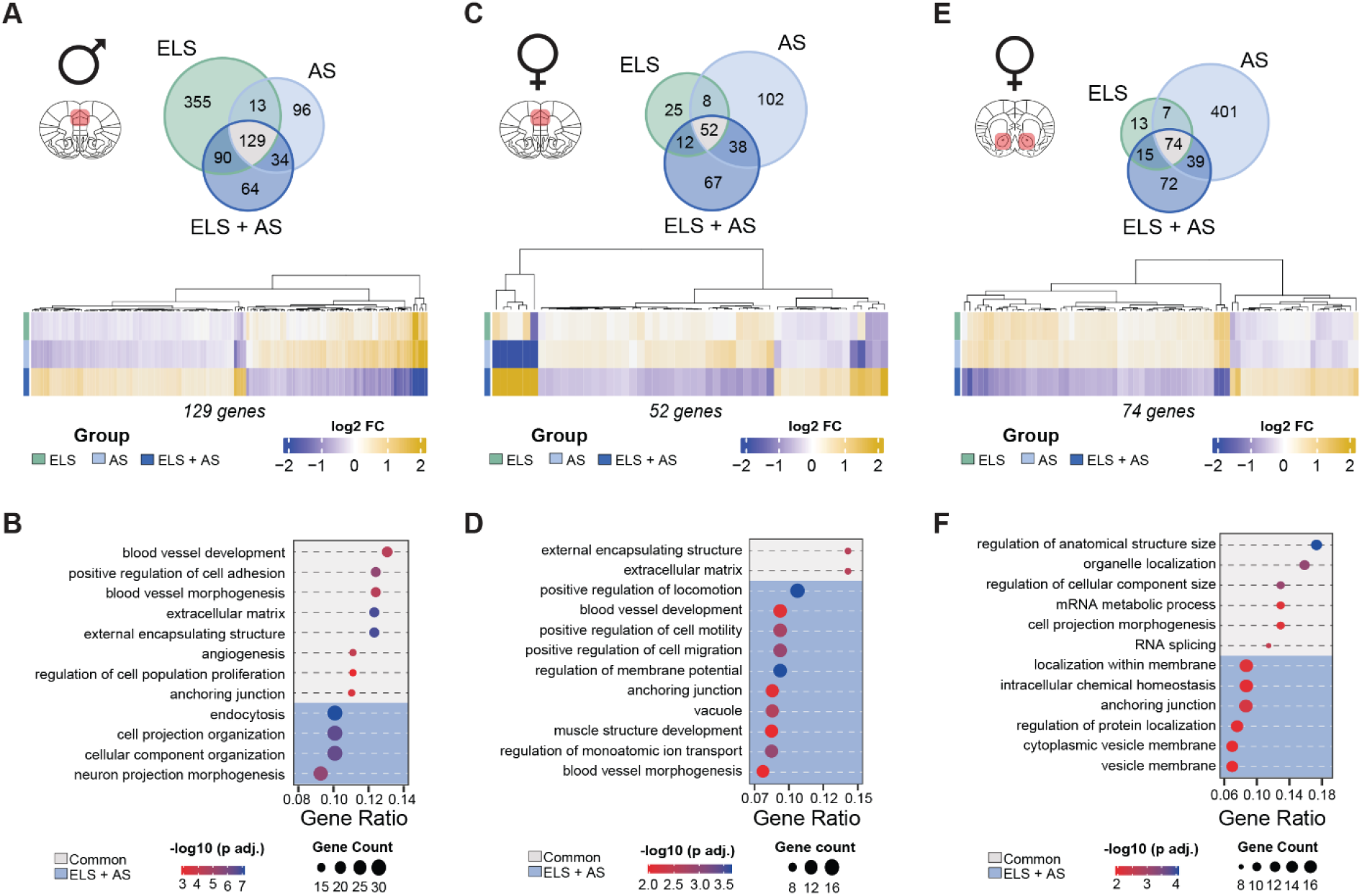
Blood-brain barrier transcriptomic alterations by ELS and AS. Differentially expressed genes in Peña et al. (2019) bulk RNA-sequencing after filtering vascular enriched genes identified by Valandewijck et al. (2018). Number of BBB regulated genes by each condition: Early-life Stress (ELS, green), Adult Chronic Stress (AS, light blue) and ELS + AS (dark blue), as well as heatmaps of the commonly regulated genes by all conditions for males PFC **(A)**, females PFC **(C)** and NAc **(E)**. Top gene ontology terms for BBB related DEGs regulated by ELS + AS (dark blue) or commonly regulated (grey) by all conditions for males PFC **(B)**, females PFC **(D)** and NAc **(F)**.

### 3.2. Early-life stress modulates susceptibility to adult stress and promotes sociability in adulthood

Early-life stress can modify substantially emotional responses and social behavior in adult life (Malave et al., 2022; Parise et al., 2025). Furthermore, ELS modulates stress responses to events occurring during adult life (Bonapersona et al., 2019; Peña et al., 2019a; Rincel et al., 2019). To better understand the long-term effects of ELS on the PFC and NAc vasculature and potential role of these alterations in stress responses, we generated and behaviorally characterized mouse cohorts exposed or not to a stressor early in life and during adulthood. First, C57Bl/6 male and female mice were subjected to 10 days of ELS by maternal separation combined with limited bedding and nesting and then, at the beginning of adult life, to 10 days of CSDS (**Fig.2A**). Control groups ran in parallel were exposed to either a single or no stressor. Social and anxiety-like behaviors were evaluated, both before and after CSDS (**Fig.2A**). ELS intervention by itself (ELS+) triggered anxiety-like behavior by decreasing the percentage of time spent in the OF center (**Fig.2B**, ELS effect: F_(1/119)_ =7.23, \**p* = 0.008) and reducing the total distance travelled in the arena (**Supp.Fig.1A**, ELS effect: F_(1/119)_ =8.18, \**p* = 0.005). Males and females displayed different profiles of exploration in the OF with ELS+ males spending less time in the center compared to ELS-females (\**p*= 0.024) and ELS+ females travelling less in the arena in comparison to unstressed ELS-females (\**p*= 0.009).

**Fig.2.**
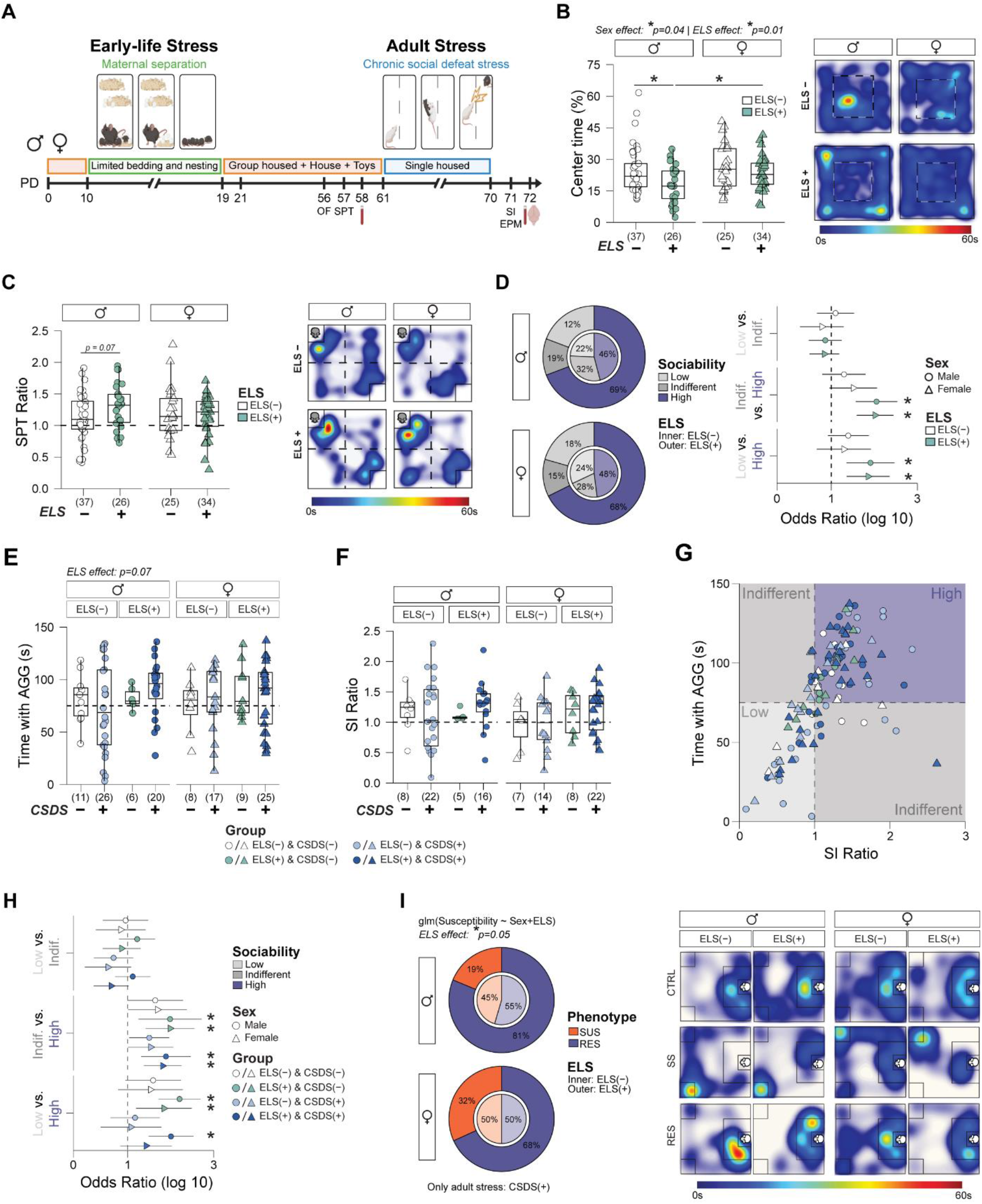
Early-life stress increases sociability behavior. Experimental timeline. Male and female mice were exposed to two-hits of chronic stress, 10-days of ELS from PD10 to 19 and a 10-days of CSDS from PD 61 to 70. Behavioral tests were performed before and after adult stress **(A)**. Long term effects of ELS revealed a reduction of time spent in the center of the Open Field (OF) **(B)** and a moderate impact on the SPT ratio for males **(C)**. Proportion of sociability profiles was increased by ELS+ and a multinomial analysis showed a likelihood of high sociability vs. low and indifferent profiles **(D)**. Social behavior after CSDS showed that ELS+ mice increased time spent with the AGG **(E)** and SI ratio **(F)**. Proportion of animals displaying a ratio higher than 1 and spent more than 50% of the trial with the AGG (purple square) **(G)**. Multinomial analysis showed that ELS increased the likelihood of high sociability profile vs low and indifferent profiles. **(H)**. A binomial logistic regression within CSDS+ animals showed that ELS+ increased proportion of resilient animals **(I)**. Data presented as boxplots with overlay of scatter plots or odds values with the 95% CI. Group size is displayed at the bottom of each figure. Representative heatmaps are presented for OF, SPT and SI. Two-way ANOVA performed for A and B. Three-way Anova performed for E and F. **\*** *p* ≤ 0.05 and tendencies ≤ 0.07 are displayed. Where applicable, multiple comparisons were corrected by Benjamini–Hochberg’s post hoc test.

ELS has been causally linked to detrimental social behavioral repertories, including but not limited to increased aggressivity or social avoidance (Cote et al., 2026; Florez et al., 2017; Parise et al., 2025; Tsuda et al., 2011). In many cases ELS followed by adult stressors, like CSDS, favor development of SUS phenotypes (Peña et al., 2019b, 2017; Rincel et al., 2019). Conversely, there is evidence suggesting that early stressful experiences can also prime positive social outcomes later in life, even after experiencing a second hit of stress (Bassey et al., 2023; Santarelli et al., 2017; Shi et al., 2021; Van Doeselaar et al., 2025). Thus, 24h after the OF test, a social preference test (SPT) was performed in the same arena (**Fig.2A**). ELS effect on sociability was not detected in variables like the SPT ratio (**Fig.2C**), time spent with the social target or social preference score (**Supp.Fig.1B, Supp.Table7**). However, the sociability classification revealed a 20% increase of males and females reaching the high sociability criteria in ELS+ groups and, an increased likelihood relative to the low (OR females: 1.60 | OR males: 1.65) or indifferent categories (OR females: 1.75 | OR males: 1.78) (**Fig.2D**). Interestingly, females travelled more during both trials of the SPT test (Trial X Sex effect: F_(1/118)_ =12.07, \**p* = 0.001) (**Supp.Fig.1C, Supp.Table7**), suggesting a sex-specific hyperlocomotion drive in social exploration. To evaluate the ELS amplifying effect on other stressors in adulthood, we chose CSDS. This paradigm produces a subpopulation of SUS mice displaying social avoidance (~2/3 of animals exposed to defeat bouts) with remaining animals defined as resilient (RES) (Golden et al., 2011; Menard et al., 2017). Regardless of the adult stress condition (CSDS), ELS+ exposed animals spent more time interacting with a novel social target (aggressor, AGG) (**Fig.2E**, ELS effect: Perm. F_(1/94)_ =3.16, *p* = 0.07) leading to higher SI ratio when compared to the ELS-/CSDS+ cohorts (**Fig.2F** ELS effect: Perm. F_(1/94)_ =3.23, *p* = 0.07). Longer time spent interacting with a novel AGG is associated with higher SI ratio, but some animals spent less than 50% of the trial interacting with the AGG potentially overshadowing ELS or CSDS effects (**Fig.2G**). Further analysis of sociability profiles revealed that exposure to ELS+ increased the percentage of high sociability (**Supp.Fig.1D**) and its likelihood relative to ELS- (OR: 4.21), low (OR females: 126 | OR males: 1.73) or indifferent animals (OR females: 1.56 | OR males: 1.63) (**Fig.2H**). In fact, exclusion of the indifferent category, confirmed that ELS+ treatment increased the percentage of RES animals, an effect modulated by sex (binomial model ELS effect with medium category *p* = 0.06, **Supp.Fig1E;** with medium category \**p* = 0.05, **Fig.2I**). Evaluation of anxiety-like behavior with the EPM did not show any effect associated to ELS or CSDS (**Supp.Fig1G, Supp.Table8**).

### 3.3. ELS in combination with CSDS modulates brain vascular-related gene expression in PFC and NAc

To further decipher the impact of ELS/CSDS on genes related to BBB integrity and function, brains of our behaviorally phenotyped mouse cohorts were collected and the expression of several genes associated to different BBB cell types (endothelial cells, mural cells and astrocytes) and stress response (glucocorticoid genes) (**Fig.3A, Fig.4A**) were evaluated by qPCR. These genes were selected for their involvement in neurovascular function, stress and mood disorders, as reported by our group and others (Cadoret et al., 2023; Dion-Albert et al., 2022a; Dudek et al., 2020; Guayasamin et al., 2025; Paton et al., 2026), and their relevance for BBB permeability, cell communication and control of stress responses (Li et al., 2022; Williams and Ghosh, 2020; Zhao et al., 2015). In male PFC, CSDS increased expression of *Fkbp5, Tjp1, Nr3c1* and *Abcc9* independently of ELS (CSDS effect: \**p* <0.05, **Supp.Table9**) with these genes clustering together (**Fig.3B**). As for ELS, it increased the expression of the pericyte marker *Atp13a5* (ELS effect: F_(1/47)_ = 3.56, *p*=0.068) and reduced expression of the angiogenic marker *Fgf2* (ELS effect: F_(1/47)_ = 3.49, *p*=0.07), this time independently of CSDS (**Fig.3C, Supp.Table9**). In female PFC, CSDS reduced expression of the angiogenic markers *Vegfa* (Phenotype effect: F_(2/45)_ =2.92, *p*=0.064) and *Fgf2* (Phenotype effect: Perm. F_(2/45)_ =2.86, *p*=0.064) in RES animals, with no effect for ELS (**Fig.3D, Supp.Table9**). The combination of ELS+ and CSDS+ stressors triggered modifications in gene expression linked to stress susceptibility or resilience (**Fig.3D-E**). For instance, for ELS+:SUS animals gene expression was reduced for the tight junction *Ocln* (\**p*=0.02 vs. ELS-:SUS and \**p*=0.02 vs. ELS+:RES) and the GC gene *Crhbp* (*p*=0.01 vs. ELS-:SUS) (**Supp.Table9**). For ELS+:RES animals, gene expression was increased for *Pdgfb*, a growth factor released by endothelial cells implicated in the communication between endothelial cells and pericytes (\**p*=0.03 vs. ELS-:RES and \**p*=0.02 vs. ELS+:SUS), and the mineralocorticoid receptor gene *Nr3c2* (\**p*=0.03 vs. ELS-:RES and \**p*=0.003 vs. ELS+:SUS) (**Fig.3E**,**Supp.Table9**). Interestingly, *Nr3c2, Ocln* and *Pdgfb* gene expression clustered together (**Fig.3D**).

**Fig.3.**
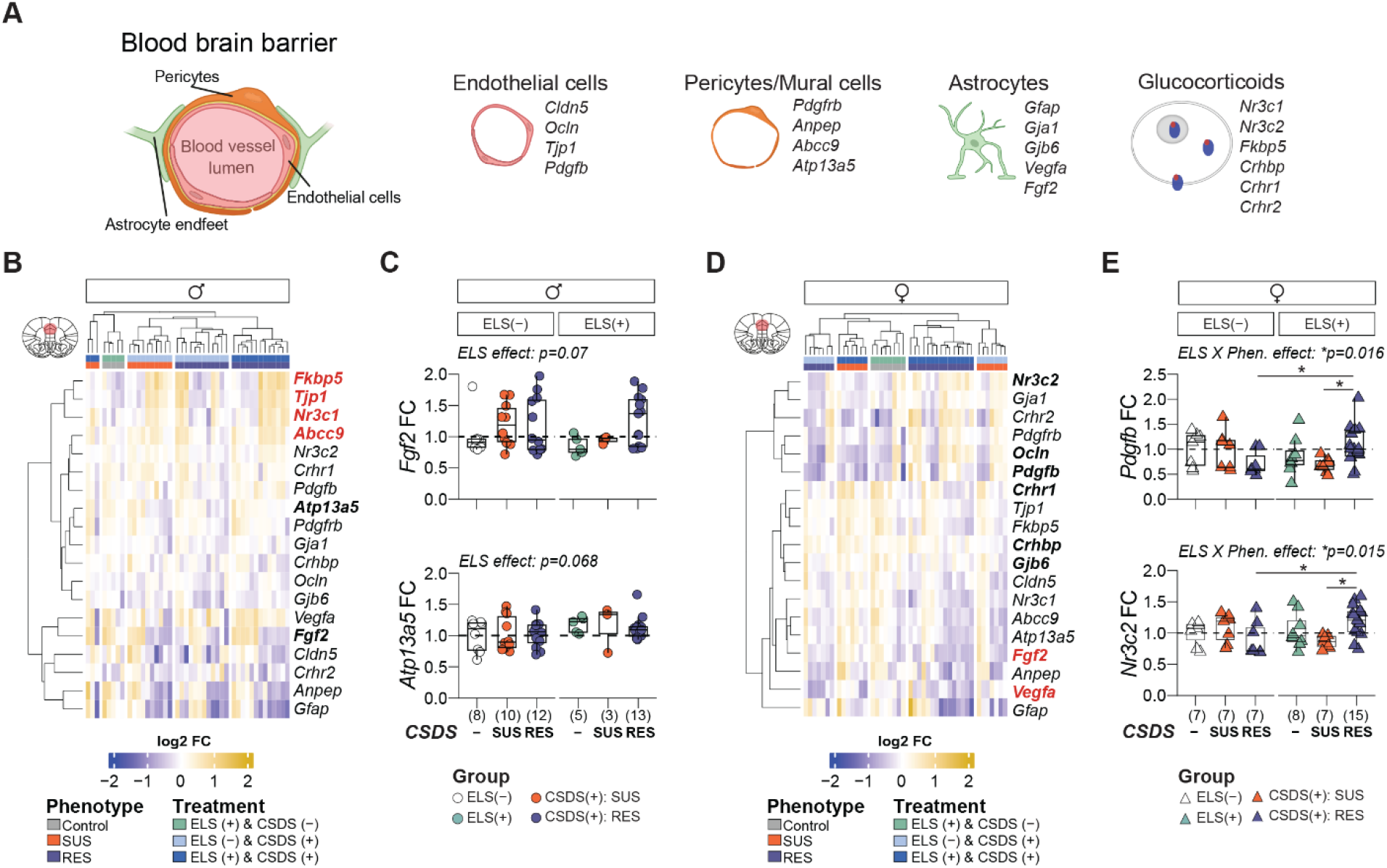
ELS and CSDS effects on PFC BBB transcriptional activity. BBB selected genes and their main associated cell type, and general glucocorticoid genes selected to evaluate the effects of ELS in combination with CSDS **(A)**. Male and female hierarchical clustering of gene expression. Group were normalized to their respective control group (ELS-:CSDS-). Gene expression presented in log2 fold change. Genes where group differences were detected are presented in bold, red for CSDS+ and black for ELS+ **(B-D)**. Fold change of representative genes modulated by ELS in males **(C)**, and two-hit condition (ELS and CSDS) in females **(E)**. Data presented as boxplots with overlay of scatter plots. Gorup size is displayed at the bottom of each figure. **\*** *p* < 0.05 and tendencies ≤ 0.07 are displayed. When apply, multiple comparisons were corrected by Benjamini–Hochberg’s post hoc test.

**Fig.4.**
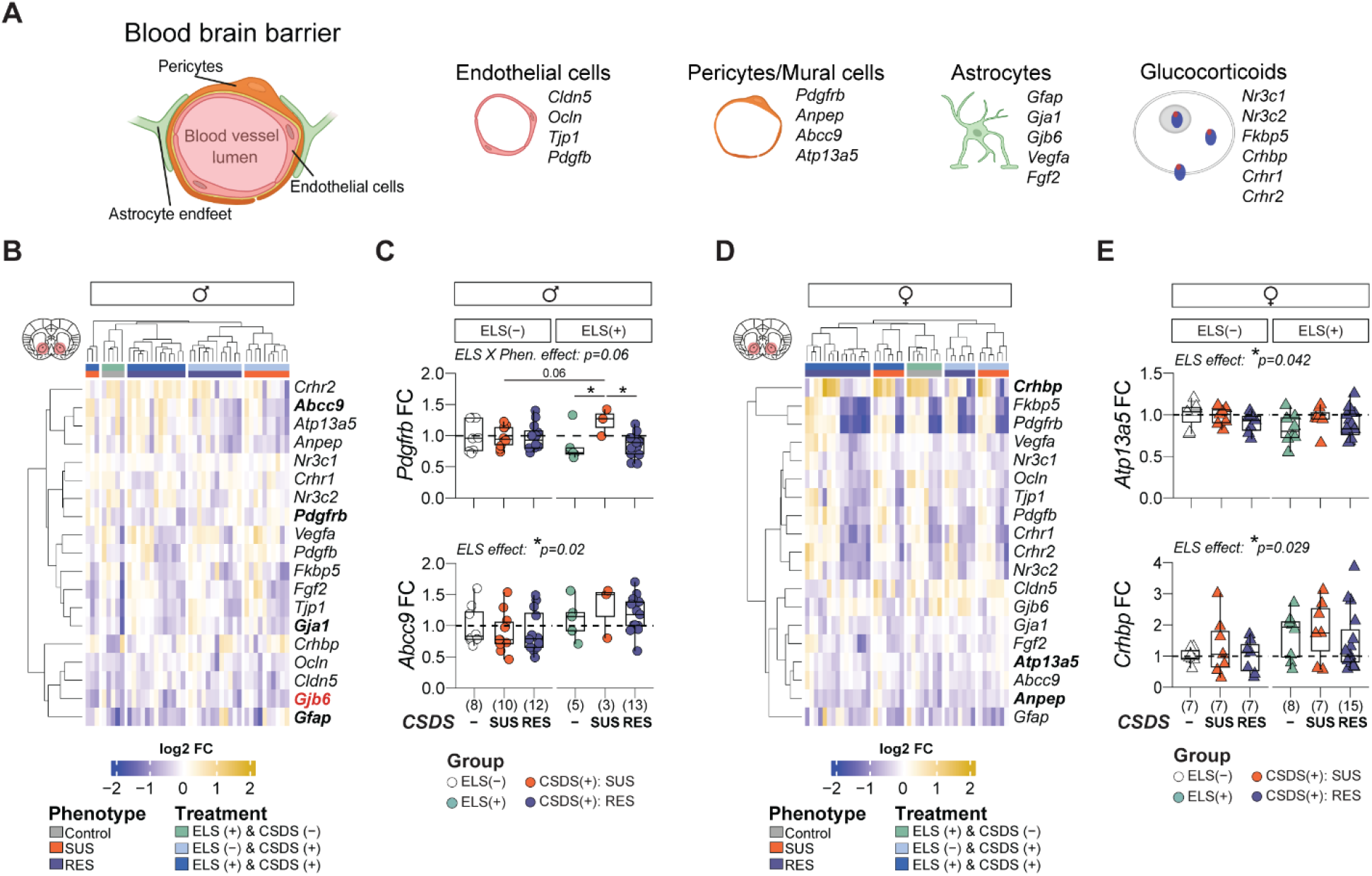
ELS and CSDS effects on NAc BBB transcriptional activity. BBB selected genes and their main associated cell type, and general glucocorticoid genes selected to evaluate the effects of ELS in combination with CSDS **(A)**. Male and female hierarchical clustering of gene expression. Group were normalized to their respective control group (ELS-:CSDS-). Gene expression presented in log2 fold change. Genes where group differences were detected are presented in bold, black for ELS+ and red for CSDS+ **(B-D)**. Fold change of representative genes modulated by the two-hit condition (ELS and CSDS) in males **(C)**, and ELS in females **(E)**. Data presented as boxplots with overlay of scatter plots. Group size is displayed at the bottom of each figure. **\*** *p* < 0.05 and tendencies ≤ 0.07 are displayed. Where applicable, multiple comparisons were corrected by Benjamini–Hochberg’s post hoc test.

In NAc, for males, combination of ELS and CSDS increased expression of *Pdgfrb*, the receptor expressed in pericytes for *Pdgfb*, specifically in ELS+:SUS animals (*p*=0.06 vs. ELS-:SUS, \**p*=0.02 vs. ELS+:CSDS-, and \**p*=0.01 vs. ELS+:RES) (**Fig.4B-C**). ELS decreased the expression of astrocytic markers *Gja1* and *Gfap*, independently of CSDS (ELS effect: \**p* <0.05, **Supp.Table10**), and increased expression of the pericyte marker *Abcc9* (ELS effect: F_(1/47)_ =3.58, *p* = 0.064, **Fig.4C**). As for CSDS, it decreased the expression of *Gjb6* in SUS animals, independently of ELS (CSDS effect: F_(1/45)_ =3.19, *p* = 0.051). Interestingly, *Gjb6* and *Gfap* expression cluster together (**Fig.4B**). For females, exposure to ELS decreased expression of *Atp13a5* and *Abcc9* pericyte markers (ELS effect: \**p* <0.05) (**Fig.4D-E**,**Supp.Table10**) and increased GC gene *Crhbp* level (ELS effect: F_(1/45)_ = 5.09, \**p* = 0.029) with all these changes independent of adult CSDS.

Together, these results show how ELS preprogram BBB-associated cellular pathways and GC signaling in adulthood in two brain areas key for regulation of mood, reward, and social behavior. Furthermore, our results suggest sex-specific neurovascular changes that can be linked to stress vulnerability and resilience.

### 3.4. ELS and CSDS desensitize HPA-axis reactivity

The GC system is highly involved in the modulation of stress responses, and the release of CORT by the adrenal glands is central for adaptative responses to environmental challenges (Lupien et al., 2009). In emotion-driven cognitive tasks, the magnitude of the CORT response varies accordingly to the intensity, number and predictability of stressful events (Koolhaas et al., 2011; Sandi, 2013). Importantly, chronicity of the stressor disrupts CORT dynamics (Kim et al., 2013; Pitman et al., 1988), and exposure to an acute stressor generally triggers a rapid elevation in CORT release, meanwhile chronic and predictable stressors appear to dampen the release of CORT (Gong et al., 2015). Additionally, GCs, in the BBB, are known to influence tight junction integrity, blood vessel stabilization, and BBB permeability (Kim et al., 2008; Kröll et al., 2009; Narang et al., 2008; Salvador et al., 2014; Williams and Ghosh, 2020). Here, to evaluate if the GC system may contribute to behavioral and neurovascular ELS-induced adaptation in the context of CSDS, CORT was measured before and after CSDS. CORT levels were reduced between pre- and post-CSDS independently of sex, ELS, or CSDS (Pre-Post effect: F_(1/83)_ =73.30, \**p* = 0.0001) (**Supp.Fig.2A, Supp.Table11**). Interestingly, after normalization of post-CSDS CORT levels on pre-CSDS, a lower reduction was observed in ELS+ exposed mice when compared to ELS-animals (ELS effect: Perm. F_(1/79)_ =4.86, \**p* = 0.003) (**Fig.5A**). Nonetheless, when controlling for the effect of ELS, the RES groups showed a smaller absolute change compared to the SUS group (*glm*, Phen. effect, \**p* = 0.045) (**Fig.5B**). Additionally, in males, increased CORT was negatively correlated with genes associated to the corticotrophin-releasing factor: *Crhbp* in the PFC (rho = −0.31, \**p* = 0.037) and *Crhr1* in NAc (rho = −0.32, \**p* = 0.034) (**Fig.5C-D**). For females, CORT increments were positively correlated with the GC chaperone gene *Fkbp5*, in both PFC (rho = 0.31, \**p* = 0.048) and NAc (rho = 0.38, \**p* = 0.012), and with *Crhr1* in NAc (rho = 0.42, \**p* = 0.006) (**Fig.5C-D**). These results suggest that ELS promotes enduring changes on GC regulation and shape adult sensitivity of the HPA-axis to a second experience of chronic stress.

**Fig.5.**
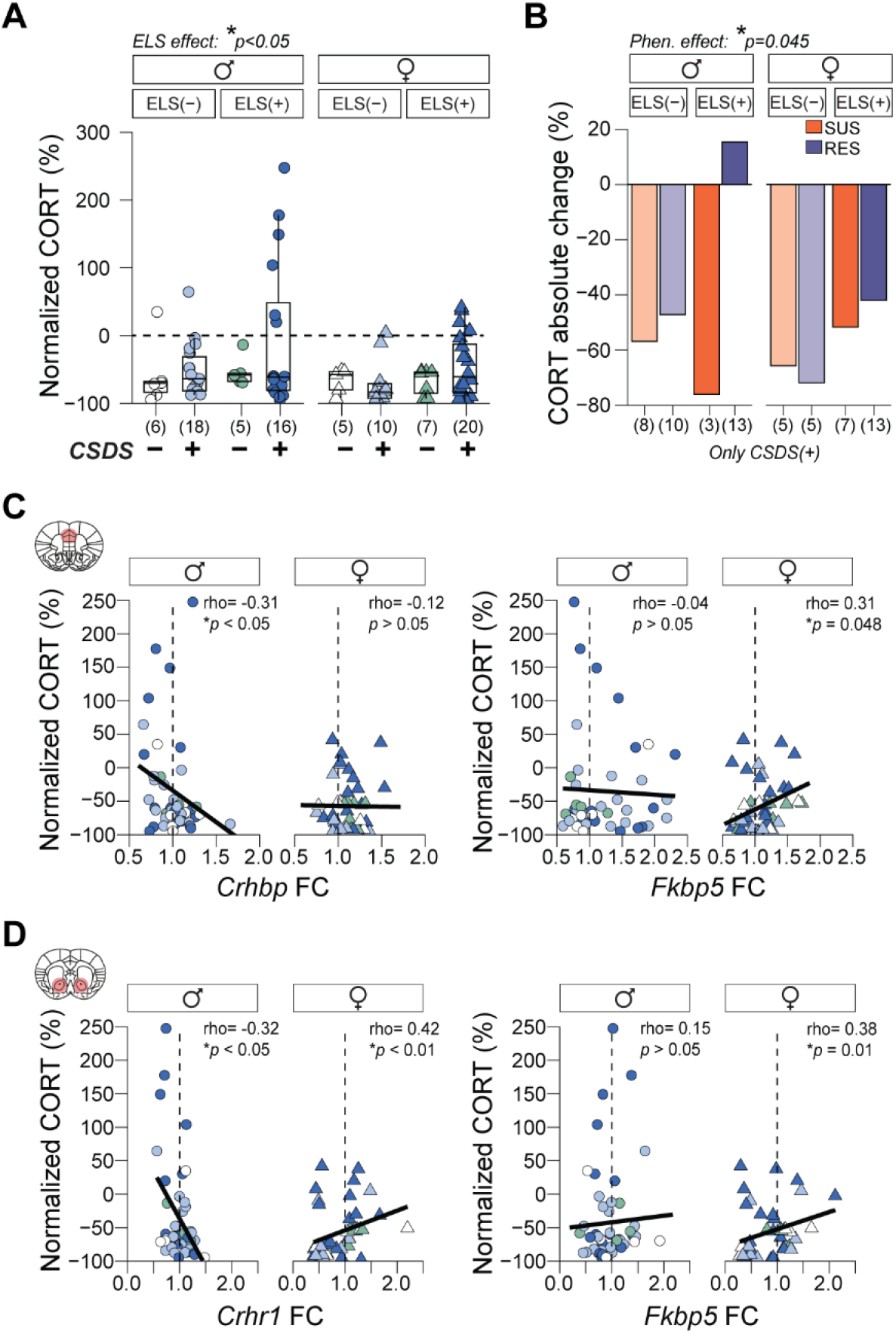
ELS and CSDS effects on NAc BBB transcriptional activity. Normalized corticosterone (CORT) expression of post-CSDS levels on pre-CSDS levels **(A)**. CORT absolute change for the groups exposed to CSDS treatment **(B)**. Spearman correlation of CORT levels with gene expression of glucocorticoid markers by sex, for both PFC **(C)** and NAc **(D)**. Data presented as boxplots with overlay of scatter plots. Group size is displayed at the bottom of each figure. **\*** *p* < 0.05.

## 4. Discussion

Structural and functional neurovascular adaptations play a critical role in shaping resilience or susceptibility to chronic stress. Dynamic modifications of the BBB, including changes in endothelial tight junction integrity (Dudek et al., 2020; Menard et al., 2017), altered communication between endothelial and mural cells (Dion-Albert et al., 2022b), and reduced astrocytic metabolic support (Dudek et al., 2025), have been implicated by our group and others in stress responses and the pathophysiology of anxio-depressive mental health disorders (Gal et al., 2023; Matsuno et al., 2022; Shi et al., 2024). Although such modifications of the BBB have been increasingly investigated in adult stress models, only a limited number of studies have examined how stress experiences during early life periods, such as childhood or adolescence, influence BBB function over short- or long-term timescales (Evertse et al., 2026; Solarz et al., 2023, 2021; Wu et al., 2022). Even less explored is how brain vasculature responds to several stressors across the lifespan. Here, our study showcases how ELS and its combination with adult stress (two-hits condition) induces BBB-specific transcriptional changes in a sex- and brain region-specific manner. We also identified a pro-social and stress resilience effect of ELS that can be associated with reduced release of CORT and upregulated gene activity in BBB cells such as pericytes or astrocytes.

ELS promotes long lasting alterations of brain transcriptional programs associated with stress reactivity and depressive-like behaviors in mice (Menezes et al., 2024; Parel et al., 2023; Parel and Peña, 2022; Peña et al., 2019b, 2017; Rincel et al., 2019). Many of those transcriptional changes are broadly conserved across brain regions but can vary according to biological factors such as sex and the presence of additional adult stress exposures (Parel and Peña, 2022; Peña et al., 2019b). For instance, in the PFC, ELS induces enduring disruptions of inhibitory signaling necessary for optimal cognitive performance in both males and females (Menezes et al., 2024). It also induces sex specific modulations of immediate early genes linked to anxiety-like responses in the female brain (Rincel et al., 2019). Intriguingly, in the NAc, transcriptional signatures from females exposed to ELS alone resemble those of human antidepressant responders, whereas combining ELS with adult stress shifts the signatures towards antidepressant nonresponders, indicating a persistent influence of ELS on mood-related circuits (Parel et al., 2023). In our analysis, BBB associated GO terms enrichment was broadly conserved across brain areas and sexes in response to stress, implicating common regulation of *blood vessels organization, cell morphogenesis*, and *vesicular organization* pathways. However, the global transcriptomic profile diverged as a function of stressor type, with ELS leading to a higher number of genes changing in the male PFC and adult stress inducing pronounced effects both in female PFC and NAc. These results pinpoint pathways through which ELS may exert persistent regulation of BBB transcriptomic profiles and influence stress responses.

Persistent ELS effects are strongly shaped by methodological parameters, like onset, type and duration of the stressor (Bonapersona et al., 2019; Murthy and Gould, 2018; Peña et al., 2019a). There is opposing evidence about long-lasting ELS effects regarding anxiety-like and social behaviors (Malave et al., 2022; Parise et al., 2025). Behavioral outcomes are frequently sexually dimorphic (Bondar et al., 2018; Cote et al., 2026; Goodwill et al., 2019) and further modulated by subsequent stress experiences across life (Peña et al., 2019a; Santarelli et al., 2017).

In our cohort, ELS exposure provoked sex-specific behavioral sensitivity, with females displaying hyperlocomotion across all behavioral tests and males showing a deeper reduction of exploratory behaviors in the OF. These results contrast with reports failing to elicit anxiety-like behavior for both sexes, while using a similar protocol as ours from PD10 to PD20 (Peña et al., 2019a, Peña et al. 2017) or using a shortened version of the protocol from PD10 to PD17 (Cote et al., 2026). Interestingly, when maternal separation was implemented from PD2 to PD14 and behavior evaluated around PD90, ELS was effective in reducing male exploration time of open and illuminated areas in the EPM (Bondar et al., 2018). Regarding social behavior in our cohort, ELS alone increased the likelihood of high sociability profile and a resilient phenotype in both males and females, contrary to previous reports where ELS increased social avoidance harnessing social preferences (Bondar et al., 2018; Cote et al., 2026; Peña et al., 2019a). Furthermore, in the two-hits condition, CSDS failed to promote the ELS-induced stress-susceptible phenotype as previously reported (Peña et al., 2019b, 2017; Rincel et al., 2019). These pieces of evidence align with the heterogeneity described in behavioral adaptation following ELS linked to methodological parameters (Bonapersona et al., 2019; Koolhaas et al., 2011; Parise et al., 2025; Sandi, 2013) and reflect how interventions with different onsets, durations, and intensities lead to specific behavioral vulnerability. They also suggest that early life stressful events with high predictability and moderate intensity can attenuate behavioral divergences and prime the development of coping strategies.

ELS promotes resilience through a shift from passive to active stress coping strategies (Cote et al., 2026; Torres-Berrío et al., 2026). Affiliative social behavior like social interactions or social buffering are active strategies to cope with stressful situations (Beery and Kaufer, 2015). Increases in affiliative behavior can be explained by environmental features surrounding stress-related manipulations, such as rearing conditions (e.g., isolation vs. group rearing) (Jaric et al., 2019; Planchez et al., 2019; Vargas et al., 2016), and enriched environment implementation during or after the stress exposure (Borba et al., 2021; Paton et al., 2026, 2023; Peña et al., 2019a). Indeed, changes in environmental experiences can reprogram the HPA-axis and alter sensitivity of the GC system to future stressors (Crofton et al., 2015; Pfau and Russo, 2015). As mentioned on Fig.2A, in our cohort, after ELS mice were group housed and had access to a house and toys. Adaptative programming of the HPA-axis after ELS has been linked to specific correlates involving the GC system, including the modulation by GCs of the chaperone protein FKBP5 in hippocampal glutamatergic neurons (Van Doeselaar et al., 2025), reduced gene expression of *Fkbp5* in the NAc (Santarelli et al., 2017), and the NR3C1 GC receptor nuclear translocation in astrocytes of the amygdala (Guayasamin et al., 2025). Consistent with a GC-dependent mechanism, our results suggest that group/environmental enrichment after ELS increased in sociability co-occurred with CORT desensitization and transcriptional changes of GC genes (*Fkbp5, Nr3c1, Nr3c2, Crhbp* and *Crhr1*).

Although bulk-tissue qPCR does not allow cell-type resolution conclusions, here by targeting genes known to be enriched in vascular cells, we were able to identify ELS neurovascular signatures associated with pericytes and astrocytes and linked them to high sociability profiles. For males, in NAc, ELS increased the expression of pericyte-related genes associated with endothelial cell communication and motility, along with decreased expression of astrocytic genes tied to endothelial cell communication. Comparable changes in pericyte-associated genes were identified in the female PFC. These profiles contrast with the RNA-seq datasets used in this study, which were associated with ELS-induced stress susceptibility. In these datasets, genes evaluated here by qPCR show ELS-related vascular alterations (*Pdgfb, Pdgfrb* and *Gja1*) in the female NAc, while CSDS-related BBB changes (*Vegfa, Pdgfb*, and *Anpep*) were more prominent in the male PFC. These differences could explain why the two cohorts - ours and Peña et al., 2019, 2017 - differ in both behavioral phenotypes and associated BBB transcriptomic profiles. Together, these patterns shed light on sex-dependent neurovascular transcriptional adaptations across PFC and NAc, reflecting a sexually dimorphic contribution of BBB pathways in response to stress. Such sex specificity reflects some of our previous reports showing sex- and brain region-specific *Cldn5* loss associated to stress susceptibility after CSDS (Dion-Albert et al., 2022b; Dudek et al., 2020; Menard et al., 2017).

Pericytes and astrocytes are fundamental, structural cell types of the BBB contributing to brain homeostasis. Pericytes are specialized mural cells controlling cerebral blood flow, vessel contractibility and vascular stabilization (Armulik et al., 2010), and astrocytes, particularly via their perivascular end-feet ensheathing blood vessels, modulate neurovascular coupling by regulation of water and ion homeostasis (Cohen-Salmon et al., 2025). Despite their central role for proper BBB and brain functions, to our knowledge there is no evidence linking pericyte or astrocyte adaptations to resilient outcomes following ELS. Notably, early life exposure to GCs has been shown to preserve brain vascular integrity and influence neurovascular stabilization by trimming vascular density, via suppression of growth factors, and by increasing pericyte coverage (Vinukonda et al., 2010).

Taken together our observations support the possibility that GC-dependent neurovascular adaptations drive stress resilience through coordinated transcriptional changes across BBB cell types. We propose that increased pericyte communication with endothelial cells, in combination with refinement of astrocyte end-feet communication with endothelial and perivascular cells, may represent a compensatory mechanism that promotes long-term BBB stabilization contributing to resilience following ELS. In this context, our results, alongside growing evidence, place pericytes and astrocytes as strategic cells integrating endocrine and immune cues shaping stress responsiveness and mood-related behaviors (Dion-Albert et al., 2022b, 2023; Dudek et al., 2025; Treccani et al., 2021). Recent findings from human postmortem tissue of individuals who encountered early life adversity support BBB cellular reorganization and alterations of transcriptional profiles of vascular cells (Wakid et al., 2026), strengthening our hypothesis of ELS-induced compensatory neurovascular adaptations, and reinforcing the need to explore further how the BBB encodes and adapts to ELS.

## Acknowledgments

This research was supported by the Canadian Institutes for Health Research (CIHR, Project Grant #427011 and #495641 to C.M.), Fonds de recherche du Quebec—Santé (FRQS, Junior 2 salary award to C.M.), C.M. Sentinel North Research Chair and a grant from Sentinel North Major Call for Proposals Phase II funded by Canada First Research Excellence Fund. C.J.P. is funded by the New York Stem Cell Foundation and NIH (R01MH129643). J.L.S. and B.D. are supported by scholarships from PhD@CERVO, NeuroQuébec, Réseau québécois sur le suicide, les troubles de l’humeur et les troubles associés (RQSHA), and Sentinel North. The authors thank all the Menard lab members for their scientific and technical contribution, especially Alice Cadoret and Luisa Bandeira Binder for assistance in sample collection, and Luisa Bandeira Binder for her help with artistic and figure aesthetics. Special thanks to the CERVO Brain Research Centre housing facility staff for their work and support.

## Author contributions

J.L.S. and C.M. designed research with scientific input from B.D., M.L. and C.J.P.; J.L.S. and B.D. performed research including behavioural experiments, molecular and biochemical analysis; J.L.S. analyzed the data and wrote the manuscript which was edited by all authors.

## Supplementary Figures

**Supp.Fig.1.**
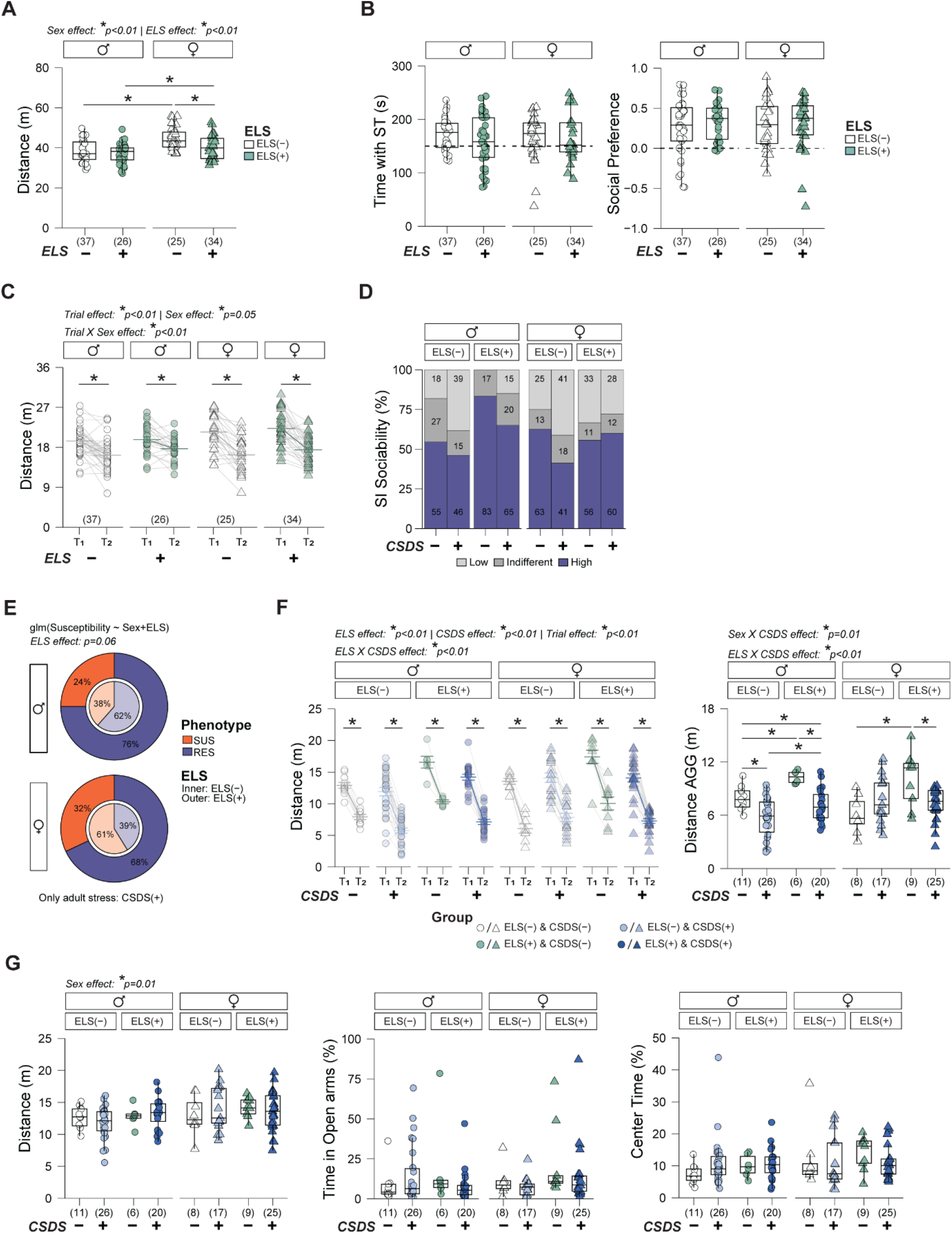
Complementary measurements of anxiety-like and social behavior. Total distance travelled in the OF **(A)**. Time with the social target during the second SPT trial and social preference index from the SPT **(B)**. Total distance travel during the two SPT trials **(C)**. Proportion of animal in each sociability category after the two-hits of stress **(D)**. A binomial logistic regression within CSDS+ animals showed a tendency of ELS+ to increase proportion of resilient animals before excluding social indifferent animals **(E)**. Total distance travel during the two SI trial (left) and a zoom in on the distance travel when the AGG was present (right) **(F)**. Anxiety-like behavior measured with the EPM made on the same day of the SI, distance travel (left), time in EPM open arms (middle) and time in EPM center (right) **(G)**. Two-way ANOVA performed for A, B and C. Three-way Anova performed for H and G. **\*** *p* ≤ 0.05 and tendencies ≤ 0.07 are displayed. Where applicable, multiple comparisons were corrected by Benjamini–Hochberg’s post hoc test.

**Supp.Fig.2.**
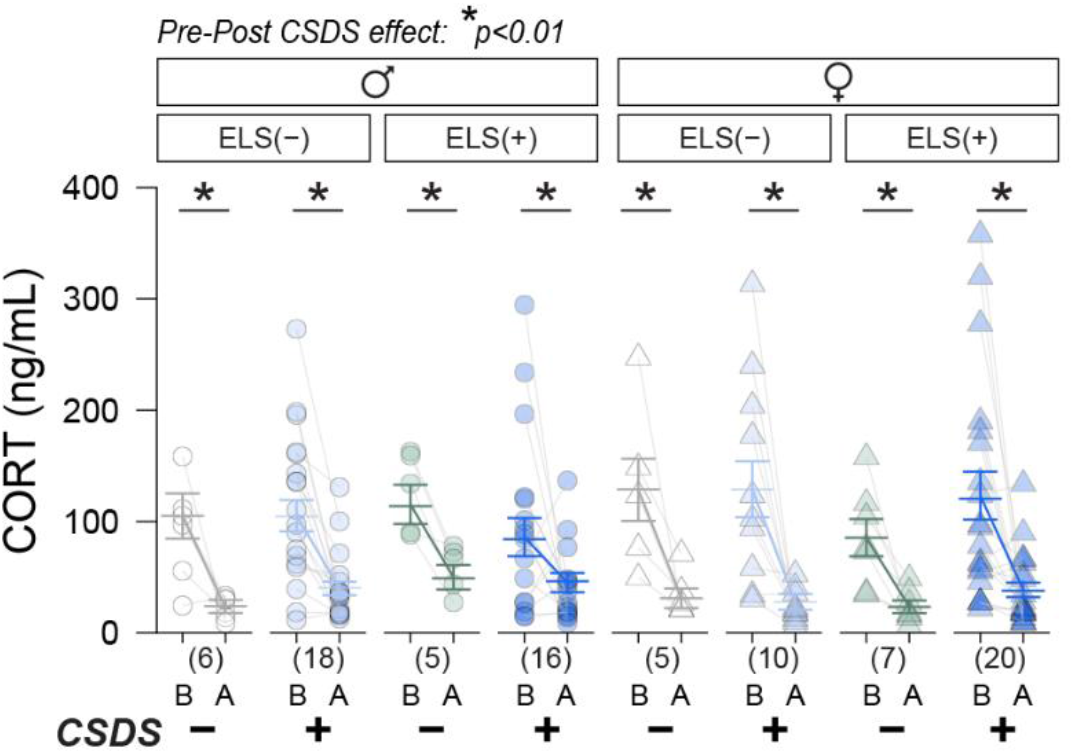
Net CORT levels before (pre-CSDS) and after (post-CSDS) adult stress in all experimental groups. Three-way Anova was performed **\*** *p* ≤ 0.05. Where applicable, multiple comparisons were corrected by Benjamini– Hochberg’s post hoc test.

